# CELEBRIMBOR: Pangenomes from metagenomes

**DOI:** 10.1101/2024.04.05.588231

**Authors:** Joel Hellewell, Samuel T. Horsfield, Johanna von Wachsmann, Tatiana Gurbich, Robert D. Finn, Zamin Iqbal, Leah W. Roberts, John A. Lees

**Author notes:** Contributed equally (Joel Hellewell, Samuel T. Horsfield).

## Abstract

**Summary:** Metagenome Assembled Genomes (MAGs) are often incomplete, with sequences missing due to errors in assembly or low coverage. Incomplete MAGs present a particular challenge for identification of shared genes within a microbial population, known as core genes, as a core gene missing in only a few assemblies will result in it being mischaracterized at a lower frequency. Here, we present CELEBRIMBOR, a snakemake pangenome analysis pipeline which uses a measure of genome completeness to automatically adjust the frequency threshold at which core genes are identified, enabling accurate core gene identification in MAGs.

**Availability and implementation:** CELEBRIMBOR is published under open source Apache 2.0 licence at https://github.com/bacpop/CELEBRIMBOR and is available as a Docker container. Supplementary material is available in the online version of the article.

## Introduction

Metagenome-assembled genomes (MAGs), generated from sequencing of complex microbial mixtures, make up a large proportion of publicly available bacterial genomes (Schmidt *et al*., 2024; Richardson *et al*., 2023), with their abundance driven by bioinformatic pipelines designed specifically to reconstruct and assay quality of MAGs (Tadrent *et al*., 2022; Kieser *et al*., 2020). A key step in the analysis of a bacterial genome collection, including for MAGs, is the identification of genes that are present in nearly all genomes for that species, known as ‘core’ genes’ (Tonkin-Hill *et al*., 2023; Zhou *et al*., 2020; Baumdicker *et al*., 2012; Parks *et al*., 2018). However, due to both uneven sequence coverage across genomes and assembly errors, MAGs are often incomplete. This results in fewer observations of genes and therefore systematic underestimation of core genome size (Li and Yin, 2022).

Existing approaches that account for incomplete genomes when estimating core genome size use binomial or multinomial models which adjust a core frequency threshold; the frequency above which a gene is assigned to the core genome (Snipen *et al*., 2009; van Tonder *et al*., 2014). Such approaches rely solely on gene frequency estimates, and so are sensitive to errors in gene prediction and clustering. PPanGGOLiN (Gautreau *et al*., 2020) uses gene synteny in addition to a multinomial model of gene frequency to adjust multiple frequency thresholds, including that of the core genome. However, PPanGGOLiN was not designed exclusively for analysis of MAGs, where systematic assembly errors may lead to a lack of contiguity, reducing identifiable synteny and therefore negatively impacting threshold adjustment.

In this work, we propose an alternative method for core frequency threshold adjustment, CELEBRIMBOR (Core ELEment Bias Removal In Metagenome Binned ORthologs). CELEBRIMBOR uses genome completeness, an estimate of how much of a genome is represented by a given assembly (Parks *et al*., 2015), jointly with gene frequency to adjust the core frequency threshold by modelling the number of gene observations with a true frequency using a Poisson binomial distribution.

## Methods

### CELEBRIMBOR workflow

CELEBRIMBOR is a snakemake workflow which conducts full pangenome analysis and core threshold adjustment. Genes are first predicted using Bakta (Schwengers *et al*., 2021) and then clustered to generate a gene presence/absence matrix using either mmseqs2 (Steinegger and Söding, 2017), or Panaroo (Tonkin-Hill *et al*., 2020), depending on user preference. Genome completeness is calculated using conserved single copy marker genes with CheckM (Parks *et al*., 2015). The presence/absence matrix and completeness are input to the core genome threshold calculation (cgt, https://github.com/bacpop/cgt), which implements the probabilistic threshold adjustment method described below.

### Core threshold adjustment

Customarily, genes are identified as core genes if they appear in 95% or more of genome samples (99% is another common choice). Due to incomplete genomes, genes with a true frequency of 95% or more can be observed in less than 95% of samples. We propose the following probabilistic model to resolve this issue.

Over *N* genome samples in a MAG dataset, the total number of times that a gene is observed is the sum of successes in *N* independently distributed Bernoulli trials that each have the probability of success *θc*_1_, …, *θc*_*N*_ where *c*_*i*_ is the completeness score of the *i*_*th*_ genome sample and *θ* is the gene frequency. The number of observations of a gene, *X*, is distributed according to a Poisson Binomial distribution. (Figure 1A).

**Figure 1:**
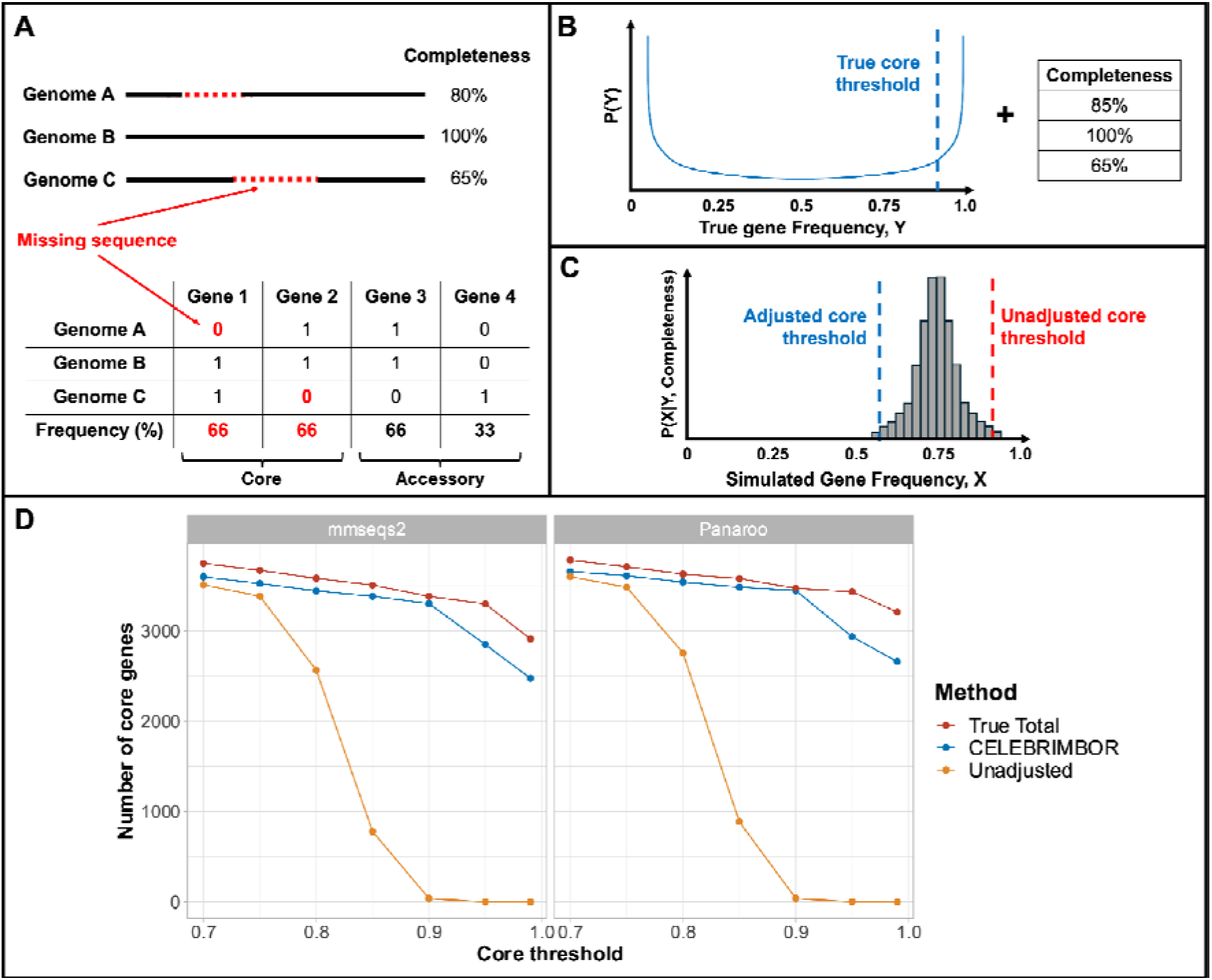
Overview of CELEBRIMBOR method and performance on simulated data. (**A**) Metagenome assembled genomes (MAGs) are often incomplete, with missing sequences from inaccurate assemblies resulting in repeated absence of gene predictions in gene presence/absence matrices. As more genomes are included in pangenome analyses, the probability of a true core gene being missed in a MAG increases, meaning that no core genes will be observed in a sufficiently large dataset. (**B**) CELEBRIMBOR uses a prior distribution of true gene frequencies, Y, given by a Beta, and completeness for each genome to simulate observed gene frequencies. (**C)** Simulated gene frequencies describe the number of times a gene is observed, X, given a true frequency Y and genome completeness. CELEBRIMBOR adjusts the core genome boundary to capture true core genes with a degree of error specified by the user. (**D**) Comparison of the number of core genes identified with varying core threshold using a simulated *E. coli* dataset, where sequences were randomly removed from assemblies. Genes with a frequency at or above the core threshold were identified as core genes. ‘True Total’ describes the number of core genes identified prior to sequence removal. ‘CELEBRIMBOR’ and ‘Unadjusted’ refer to the CELEBRIMBOR-adjusted and unadjusted estimates of the number of core genes respectively. Genes were clustered using mmseqs2 and Panaroo.

The true frequencies of the genes in the pangenome are assumed to have a U-shaped Beta distribution (Lobkovsky *et al*., 2013; Colquhoun *et al*., 2021), where most genes are either core genes with a high true frequency or are much rarer accessory genes (Figure 1B).

For a core gene *G* with unknown frequency *θ* >= 0.95, we estimate through simulations the probability of observing this gene *X* or fewer times over *N* genome samples. We then select the value of *X* such that this probability is 5% (by default, but this value is tunable) or less: *P*(*observe G* <= *X times* | *θ* >= 0.95) <= 5% . (Figure 1C).

This chosen value of *X* is the new, adjusted core threshold that is used to identify core genes in the MAG data. For any new threshold, there will be a corresponding probability that a gene with a true unknown frequency less than 95% will appear *X* or more times. This is the probability of wrongly identifying a non-core gene as core. By default, CELEBRIMBOR prioritises maximising the probability of identifying true core genes over minimising the probability of wrongly identifying non-core genes as core.

### Simulated MAG dataset

*Escherichia coli* read datasets were randomly selected from (Kallonen *et al*., 2017) using SPAdes assemblies from (Page *et al*., 2016). Random sections of each genome were removed using a custom script (https://github.com/bacpop/MAG_pangenome_pipeline/blob/main/simulate_pangenomes/remove_sequence.py), which removes a proportion of a genome assembly based on previously observed MAG completeness values from MGnify (Richardson *et al*., 2023).

### Pangenome analysis

Assemblies pre- and post-sequence removal were analysed using CELEBRIMBOR, using Bakta v1.8.2 (Schwengers *et al*., 2021) for gene prediction and annotation, mmseqs2 v14.7e284 (Steinegger and Söding, 2017) or Panaroo v1.4.1 (Tonkin-Hill *et al*., 2020) in strict mode for clustering, and CheckM v1.2.2 (Parks *et al*., 2015) for genome completeness estimation. CELEBRIMBOR v0.1.0 and PPanGGOLiN v2.0.3 (Gautreau *et al*., 2020) were run on both pre- and post-sequence removal dataset. CELEBRIMBOR was run using core gene frequency thresholds in the range 70%-99%, with rare gene frequency threshold and error rate set at 5% for all analyses. PPanGGOLiN was run with three pangenome partitions (‘-K 3’) on GFF files generated by CELEBRIMBOR via Bakta.

The number of genes labelled as ‘core’ and ‘rare’ by CELEBRIMBOR, or ‘persistent’ and ‘cloud’ by PPanGGOLiN were compared between pre- and post-sequence removal datasets across the range of core thresholds specified above. A full description of the simulation workflow used for CELEBRIMBOR benchmarking is available on Github (https://github.com/bacpop/MAG_pangenome_pipeline/tree/main/simulate_pangenomes). All analysis was performed on a workstation with 2 × 20 core Intel Xeon Gold CPUs.

### Code availability

Code for CELEBRIMBOR and pangenome simulations is available on Github (https://github.com/bacpop/CELEBRIMBOR). Code for cgt is also available on Github (https://github.com/bacpop/cgt).

## Results

To simulate the effect of a dataset containing incomplete assemblies on core genome size estimation, we randomly removed sequence blocks from 500 full *Escherichia coli* genome assemblies (Kallonen *et al*., 2017), with the amount of sequence removed from each assembly guided by previously observed MAG completeness values (Richardson *et al*., 2023). The dataset pre- and post-sequence removal was analysed using CELEBRIMBOR using two different methods of gene clustering; mmseqs2, a highly scalable protein sequence clustering method (Steinegger and Söding, 2017), and Panaroo, a start-of-the-art pangenome analysis method which uses gene frequency and synteny to cluster and correct gene prediction errors (Tonkin-Hill *et al*., 2020).

As the core threshold is increased past 75%, the number of core genes identified falls dramatically, reaching zero at 90% without threshold adjustment for both mmseqs2 and Panaroo (Figure 1D). Using CELEBRIMBOR greatly improves estimates of core genome size, staying close to the true total of core genes, independent of clustering method.

Both clustering methods gave similar core genome size estimates after correction by cgt; at a core threshold of 95%, mmseqs2 and Panaroo estimated a total of 3299 and 3434 core genes pre-sequence removal, and 2850 and 2936 using CELEBRIMBOR’s post-sequence removal, whilst only 2 and 3 core genes were identified without adjustment respectively (Supplementary Table 1). The slightly larger core genome estimates provided by Panaroo over mmseqs2 is indicative of the more accurate orthologue detection strategies employed by Panaroo, including use of iterative similarity thresholds and synteny during clustering. Altering Panaroo’s stringency settings did not affect estimates of core genome size, although more stringent removal of low frequency genes reduced the number of estimated rare genes (Supplementary Figure 1). In comparison, PPanGGOLiN estimated a total of 3711 and 4099 core genes pre- and post-sequence removal respectively. A higher core genome size estimate post-sequence removal is indicative of incorrect assignment of accessory genes as core, meaning PPanGGOLiN’s method for threshold adjustment is overly sensitive and results in false positives. CELEBRIMBOR is by default more conservative, resulting in more false negatives, although this can be reduced by increasing the threshold error value.

To compare computational efficiency, a series of datasets up to 1500 simulated MAGs in size from (Kallonen *et al*., 2017) were analysed using CELEBRIMBOR running mmseq2 or Panaroo. Using mmseqs2 gave consistently lower runtimes compared to Panaroo, although runtime for both workflows scaled linearly with dataset size (Supplementary Figure 2). Both workflows used a maximum of 35 Gb memory for all analyses, as this was the pre-run memory allocation designated by snakemake. Overall, mmseqs2 provides greater computational scalability in clustering over Panaroo, which is of particular importance when including increasingly large numbers of genomes in analyses.

## Discussion

MAGs are a crucial resource in the analysis of bacterial pangenome diversity, particularly for species that cannot be cultured under artificial conditions. CELEBRIMBOR enables a parametric recapitulation of the core genome using MAGs, which would otherwise be unidentifiable due to missing sequences resulting from errors in the assembly process. CELEBRIMBOR implements both computational efficient and accurate clustering workflows; mmseqs2, which scales to millions of gene sequences (Steinegger and Söding, 2017), and Panaroo, which uses sophisticated network-based approaches to correct errors in gene prediction and clustering (Tonkin-Hill *et al*., 2020).

We show that CELEBRIMBOR can accurately estimate the size of the core genome in simulated MAG data, far outperforming methods where no threshold adjustment is made. Panaroo and mmseqs2 were not able to identify any core genes past the 90% threshold, despite Panaroo including a method for “re-finding”, which identifies missing gene predictions resulting from mutations or assembly errors (Tonkin-Hill *et al*., 2020). However, integration of Panaroo into CELEBRIMBOR combines Panaroo’s error correction methods with CELEBRIMBOR’s core threshold adjustment, resulting in highly accurate quantification of gene frequencies of both core and lower frequency genes.

A crucial assumption underlying core genome threshold adjustment models (including in this work) is that genes in the genomes in the MAGs are assumed to be missing completely at random. This entails that the causal mechanisms that are responsible for the missing data are independent of the data. However, if certain parts of the genome are more liable to errors in the steps before core gene threshold adjustment (sequencing, assembly, labelling, etc) because of features of the genomic data, then this violates the missing completely at random assumption. A likely consequence of this is that true core genes that are not missing at random could fail to be labelled as core genes. Therefore, CELEBRIMBOR may fail to correctly identify core genes in regions that are systematically misassembled.

CELEBRIMBOR is a rapid pangenome analysis and core threshold adjustment pipeline designed for analysis of large MAG datasets. It enables researchers working exclusively with MAGs to accurately identify core genes, enabling epidemiological and evolutionary analysis even in the presence of missing data, as well as investigation of sequencing and assembly issues that lead to gene drop out in gene presence/absence matrices.

