## Supplementary material for "CELEBRIMBOR: Pangenomes from metagenomes"

### Supplementary Figures

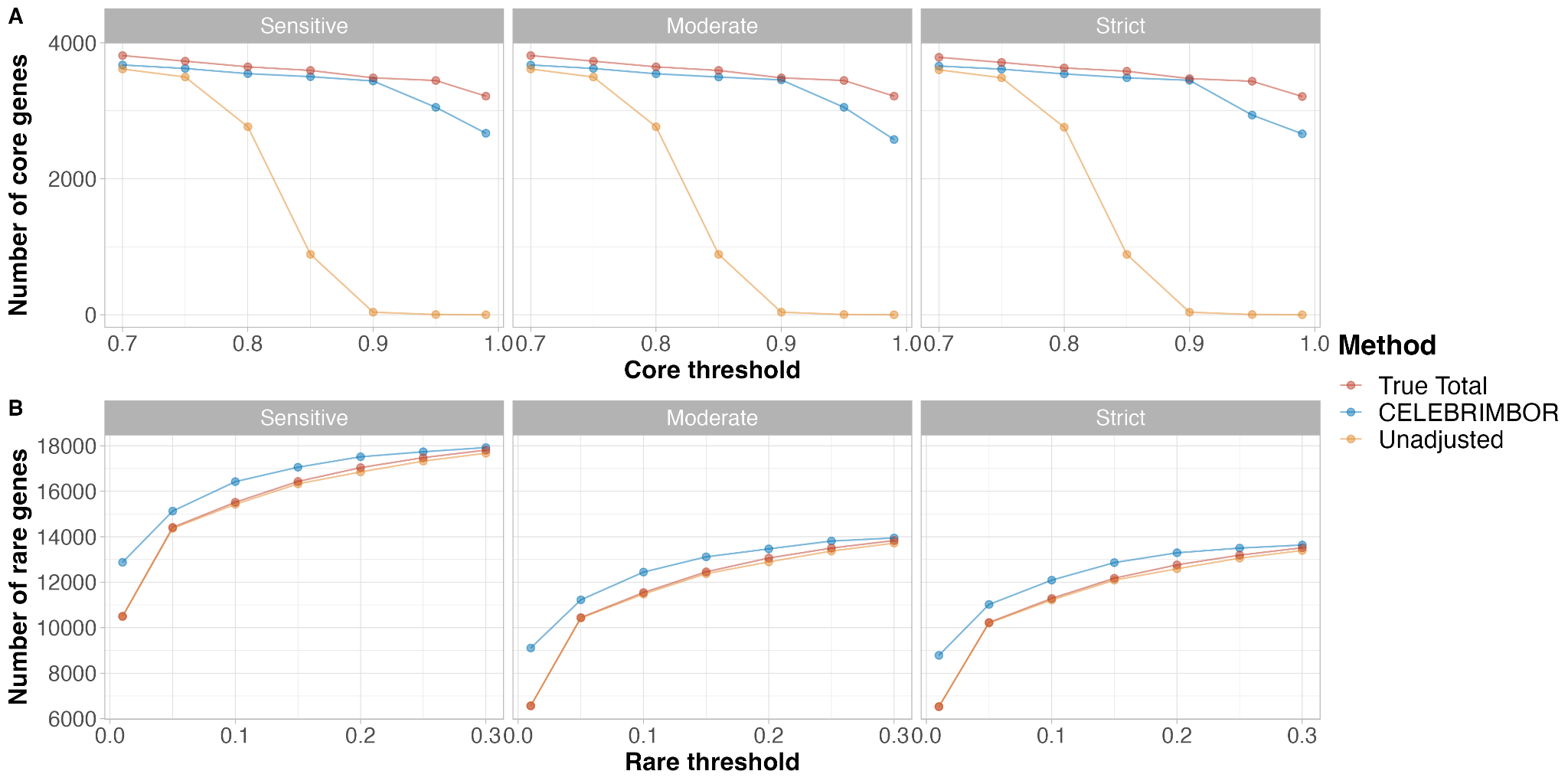

Supplementary Figure 1: Comparison of Panaroo stringency settings on estimated number of (**A**) core and (**B**) rare genes using CELEBRIMBOR. For (**A**), the rare threshold was set at 5%; for (**B**) the core threshold was set at 95%. For both (**A**) and (**B**), the error threshold was set at 5%. Columns describe Panaroo stringency settings; ‘Sensitive’, ‘Moderate’ and ‘Strict’ (see (Tonkin-Hill *et al.*, 2020) for description of settings).

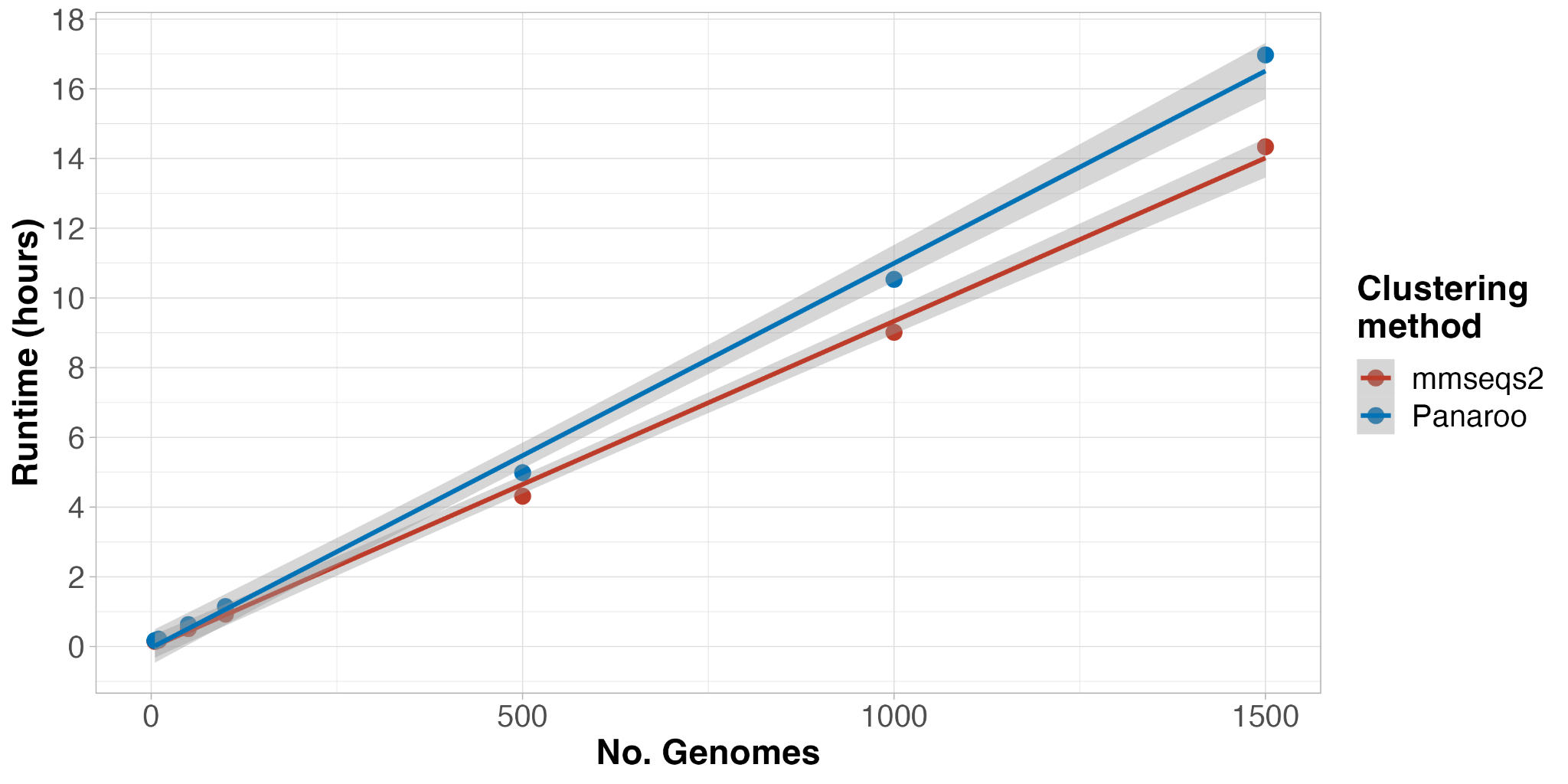

Supplementary Figure 2: Runtime comparison of clustering methods used in CELEBRIMBOR. Grey boundaries indicate 95% confidence intervals for linear regression. Comparisons were conducted on simulated MAGs from *E. coli* genomes from (Kallonen *et al.*, 2017). CELEBRIMBOR was run with 16 threads for all analyses.

### Supplementary Tables

Supplementary Table 1: Number of genes assigned to frequency compartments by different clustering algorithms. For clustering using mmseqs2 and Panaroo, CELEBRIMBOR was run with core and rare thresholds set to 95% and 5% respectively, and error threshold at 5%. Intermediate genes are found at frequency 5% > X > 95%.

| **Clustering method** | **Adjustment** | **No. core** | **No. intermediate** | **No. rare** |
| --- | --- | --- | --- | --- |
| mmseqs2 | True | 3299 | 4929 | 24726 |
|  | CELEBRIMBOR | 2850 | 3973 | 26131 |
|  | Unadjusted | 2 | 7791 | 25161 |
| Panaroo | True | 3434 | 4646 | 10232 |
|  | CELEBRIMBOR | 2936 | 3989 | 11022 |
|  | Unadjusted | 3 | 7739 | 10205 |
| Ppanggolin | True | 3711 | 2955 | 15149 |
|  | Adjusted | 4099 | 2824 | 14453 |
